# Doublecortin-like kinase 1 is a therapeutic target in squamous cell carcinoma

**DOI:** 10.1101/2022.05.26.493670

**Authors:** David Standing, Levi Arnold, Prasad Dandawate, Brendan Ottemann, Vusala Snyder, Sivapriya Ponnurangam, Afreen Sayed, Dharmalingam Subramaniam, Pugazhendhi Srinivasan, Sonali Choudhury, Jacob New, Deep Kwatra, Prabhu Ramamoorthy, Badal C. Roy, Melissa Shadoin, Raed Al-Rajabi, Maura O’Neil, Sumedha Gunewardena, John Ashcraft, Shahid Umar, Scott J. Weir, Ossama Tawfik, Subhash B. Padhye, Shrikant Anant, Sufi Mary Thomas

## Abstract

Doublecortin like kinase 1 (DCLK1) plays a crucial role in several cancers including colon and pancreatic adenocarcinomas. However, its role in squamous cell carcinoma (SCC) remains unknown. To this end, we examined DCLK1 expression in head and neck squamous cell carcinoma (HNSCC) and anal squamous cell carcinoma (ASCC). We found that DCLK1 is elevated in patient SCC tissue, which correlated with cancer progression and poorer overall survival. Furthermore, DCLK1 expression is significantly elevated in HPV negative cancer tissues, which are typically aggressive with poor responses to radiation therapy. To understand the role of DCLK1 in tumorigenesis, we used specific shRNA to suppress DCLK1 expression. This significantly reduced tumor growth, spheroid formation, and migration of HNSCC cancer cells. To further the translational relevance of our studies, we sought to identify a selective DCLK1 inhibitor. Current attempts to target DCLK1 using pharmacologic approaches have relied on non-specific suppression of DCLK1 kinase activity. Here, we demonstrate that DiFiD [3,5-bis (2,4-difluorobenzylidene)-4-piperidone] binds to DCLK1 with high selectivity. Moreover, DiFiD mediated suppression of DCLK1 led to G2/M arrest and apoptosis and significantly suppressed tumor growth of HNSCC xenografts and ASCC patient derived xenografts, supporting that DCLK1 is critical for SCC growth.

## Introduction

Squamous cell carcinoma (SCC) originates in the mucosal epithelium at several sites including the oral cavity, oropharynx, and anus. Head and neck squamous cell carcinoma (HNSCC) is the sixth most common cancer worldwide(1). Etiologic factors associated with HNSCC include tobacco and alcohol abuse, and human papilloma virus (HPV) infection. Despite therapeutic advances, HNSCC remains difficult to treat; due, in part, to late-stage tumor presentation, as well as high rates of local recurrence and distant metastases that contribute to poor five-year survival (2, 3). Anal squamous cell carcinoma (ASCC) is a rare cancer, which like HNSCC is associated with HPV infection, and its incidence is increasing. Accordingly, the identification of novel targets and therapeutics is needed to improve SCC treatment outcomes. There remains a paucity of preclinical descriptions of ASCC biology or potential drug targets.

Recently, doublecortin like kinase 1 (DCLK1) was found to be highly expressed in salivary gland tumors and is associated with low overall and disease-free survival (4). DCLK1, encodes a member of the doublecortin family and protein kinase superfamily (5, 6). DCLK1 contains two N-terminal doublecortin domains and a C-terminal kinase domain homologous to Ca^2+^/calmodulin-dependent kinases. Doublecortin domains participate in microtubule polymerization, whereas the serine/threonine protein kinase domain and a serine/proline-rich domain, mediate multiple protein-protein interactions. The activity of each domain is independent of one another (7). DCLK1 is expressed in various cell types including neurons, osteoblasts, and tuft cells in the colon. Initial reports by our group (5, 6, 8, 9) and that of multiple laboratories (10–12), have reported DCLK1 expression is observed in a small percentage of cells (<5%) in colon and pancreatic adenocarcinomas. Further, it has been established as a marker of tumor-initiating cells (13). The role of DCLK1 in SCCs remains unknown.

Presently, we determine that DCLK1 expression is upregulated in SCC. Further, DCLK1 regulates HNSCC proliferation, invasion, and migration. We also report for the first time that DiFiD [3,5-bis (2,4-difluorobenzylidene)-4-piperidone], which has shown efficacy in preclinical models of pancreatic cancer, binds to and inhibits DCLK1 activity and demonstrated antitumor effects in SCC (14). These studies demonstrate the potential therapeutic value of targeting DCLK1 in SCC.

## Methods and Materials

### Cell lines, patient tissues, and reagents

Well-characterized HNSCC cell lines UM-SCC-1 (from Dr. Tom Carey, University of Michigan, Ann Arbor, MI), OSC19 (from Theresa Whiteside, University of Pittsburgh, Pittsburgh, PA), HN5 (from Dr. Jeffrey Myers, The University of Texas MD Anderson Cancer Center, Houston, TX) and FaDu (ATCC) were used in this study (15). Het-1A, an immortalized non-cancerous esophageal squamous epithelial cell line was obtained from ATCC. Established cell lines were authenticated by short tandem repeat profiling at Johns Hopkins in 2018 using the Promega GenePrint 10 kit and analyzed using GeneMapper v4.0 software. FaDu were maintained in EMEM (Corning) with 10% heat inactivated FBS. All other cell lines were maintained in DMEM (Corning) with 10% heat-inactivated FBS (Sigma-Aldrich) without antibiotics. Cells were incubated at 37°C in the presence of 5% CO_2_. Tissue Microarray (#HN483) was purchased from US Biomax, Inc. Sections from paraffin embedded blocks of seventeen deidentified anal squamous cell carcinoma (ASCC) patient samples were obtained from the Department of Pathology, University of Kansas Medical Center. After review, the institutional IRB determined that this study (STUDY00140502) was not research involving human subjects. DiFiD and EF24 were acquired as previously reported (14, 16, 17). KN-62 was purchased from Tocris Bioscience (Minneapolis, MN).

### Apoptosis assays

For apoptosis, Apo-one Homogeneous Caspase-3/7 Assay kit was performed according to the manufacturer’s protocol to calculate caspase 3/7 activity (Promega Corporation, Madison, WI). Additionally, Annexin V/PI staining was performed. In short, 1 х 10^5^ cells were plated in 10 cm dishes and allowed to grow for 24 hours, after which they were treated with DMSO or DiFiD at IC_50_ doses. Following treatment for 48 hours, cells were washed with phosphate buffered saline (PBS), collected, and stained with Annexin V antibody conjugated with a FITC fluorophore and propidium iodide (PI). Stained cells were then assessed by flow cytometry.

### cDNA synthesis and Real-time qPCR

FaDu (2x10^5^) cells were seeded in 6-well tissue culture dishes. shRNA clones were maintained as above. Total RNA was isolated with Trizol reagent. cDNA was generated from the total RNA by using Maxima H Minus Reverse Transcriptase (200 U/L) according to manufacturer’s protocol (Thermo Fisher Scientific). Quantitative Real-time PCR (qPCR) was carried out with SsoAdvanced Universal SYBR Green Supermix (BioRad) according to the manufacturer’s protocol. Crossing threshold values for genes were normalized to β-actin. Changes in mRNA expression are represented as fold change relative to control. Primers used in this study were as follows: β-actin: 5′-GCTGATCCACATCTGCTGG-3′ and 5′-ATCATTGCTCCTCCTCAGCG-3′; Hes1: 5’-GGTCAGTCACTTAATACAGCTCTCTC-3’ and 5’-CCTCTCTTCCCTCCGGACT-3’; DCLK1: 5’-AGGCCATCATTAACCCAG-3’ and 5’-AGGTGGACTTTCCTTCTCCATAC-3’; ALDH1A1: 5’-AGCAGGAGTGTTTACCAAAGA-3’ and 5’-CCCAGTTCTCTTCCATTTCCAG-3’.

### Cell cycle analysis

HNSCC cells (HN5 and FaDu) were seeded at 1x10^5^ cells in 100 mm tissue culture dishes for 24h. Cells were then treated and incubated an additional 24 or 48 hours. Cells were then trypsinized and suspended in PBS. Single-cell suspensions were fixed using 70% ethanol in PBS for 2 hours. The cells were permeabilized with PBS containing 1 mg/ml propidium iodide (Sigma-Aldrich), 0.1% Triton X-100 (Sigma-Aldrich) and 2 µg DNase-free RNase (Sigma- Aldrich) at room temperature. Flow cytometry was performed with a BD LSRII (Becton Dickinson, Mountain, View, CA), capturing 10,000 events for each sample. Results were analyzed with WEASEL software (Chromocyte, Electric Works, Sheffield, UK).

### Cloning of DCLK1 DCX Domain

DCLK1 DCX domain was amplified by PCR. cDNA was run on 2% agarose gels and bands were purified using the GeneJET gel purification kit (Thermo Fisher Scientific, Waltham, MA) according to the manufacturer’s protocol. DCX DNA was cloned into the N-Terminal pFLAG 3 vector (Addgene) by restriction digest using NotI and BamHI restriction enzymes. FaDu cells were transfected and selected using 400 µg/mL G418 sulfate salt. Primers used for DCX Domain:

DCLK1 NotI FP: 5’-GATCGCGGCCGCGATGTCCTTCGGCAGA-3’ DCLK1 BamHI DCX RP: 5’-GATCGGATCCCTAGCCATCGTTCTCATC-3’

### DCLK1 Kinase assay

Purified recombinant DCLK1, CamKIIα, CamKIIβ and CamKIV (150 ng, Signalchem) were incubated in reaction buffer (Invitrogen) with 9 nM peptide substrate (Gs peptide), 10 µM ATP (Invitrogen), and either DMSO, EF24, DiFiD or KN-62 (a pan-CAMK inhibitor) for 30 min at 37°C. We used Gs peptide because it has been used as a substrate for CamK assays (18, 19). Our assays were adapted based on these studies. Fold change was calculated relative to ATP counts in the control. Consequently, 100 μl of luciferin-luciferase mixture (ATP determination kit, Invitrogen) was added into each well and luminance was monitored at 560 nm by using Synergy^TM^ NEO microplate reader.

### Drug affinity responsive target stability (DARTS)

The ability of DiFiD to interact with and stabilize DCLK1 in cells was studied by using DARTS (20). FaDu cells were cultured and grown up to 70-80% confluency. Cells were washed three times with ice-cold PBS and. lysed using M-PER lysis buffer supplemented with protease inhibitor cocktail tablet (Roche) and collected by scraping off with a cell scraper. Cells were lysed on ice for 10 minutes, clarified by centrifugation, and protein estimated using BCA method. Cell lysates were divided into equal concentration aliquots and incubated with DMSO or DiFiD (5 μM) for 30 minutes with gentle shaking. Following incubation, lysates were treated with pronase (10 mg/ml stock) in 1:1/100, 1:1/200, 1:1/400, 1:1/800, 1:1/1600, 1:1/3200, 1:1/6400 protein:pronase ratio for 15 minutes. The protease digestion was stopped using 20X protease inhibitor cocktail (Roche) for 10 minutes on ice. The resultant protein samples were diluted with 4X Laemmli buffer, heated at 70°C for 10 minutes and loaded on to 10% SDS-PAGE gel, transferred to PVDF membrane and incubated with DCLK1 antibody at a concentration of 1:1000. Protein levels on western blot were pictured by Bio-Rad ChemiDoc-XRS+ instrument and analyzed by image lab software. Band intensity was calculated relative to the lowest dilution of pronase.

### Immunoblotting

HNSCC cells (HN5, UMSCC1, OSC19, FaDu; 1x10^6^ cells) were seeded in 100 mm tissue culture dishes. Cells were treated with respective IC_50_ µM concentrations of DiFiD or vehicle control for 24 and 48 hours. Fifty µg of the total protein lysate were subjected to polyacrylamide electrophoresis and transferred to PVDF membranes with a wet-blot transfer apparatus (Bio- Rad, Hercules, CA). After blocking in 5% milk overnight and incubation with primary antibody (using manufacturer recommended dilutions), the signal was developed with horseradish peroxidase-conjugated secondary antibody (dilution 1:5000) and ECL Western blotting detection reagents (Amersham-Pharmacia, Piscataway, NJ). Actin and GAPDH were used as loading controls. Actin (#A1978) and DCLK1 (targeted to the N-terminus, #62257) (#SAB4200186) were purchased from Sigma Aldrich (Sigma-Aldrich, St. Louis, MO). Bax (#5023), PARP (#9542), Cyclin B1 (#12231), CyclinD1 (#2978), CDC2 (#77055) were purchased from Cell Signaling Technologies (Beverly, MA). GAPDH (#SC-47724) was purchased from Santa Cruz Biotechnology (Dallas, TX).

### Immunohistochemistry

Paraffin-embedded tissues were cut to 4 μm sections, deparaffinized and subjected to antigen retrieval. The tissue sections were blocked with UltraVision Hydrogen Peroxide block for 10 minutes (Thermo Scientific). The slides were incubated with primary antibodies (1:200 dilution) overnight at 4^°^C. DCLK1 antibody was purchased from Abcam (#37994) (Cambridge, MA) and used according to a previously published protocol (21). The following day, the primary antibody was washed, and tissues were incubated with HRP Polymer Quanto for 10 minutes then developed with a DAB Quanto Chromogen-Substrate mixture and counterstained with hematoxylin. The slides were assessed using a Nikon Eclipse Ti microscope under a 200X magnification. Staining intensity was determined by Aperio Imaging Software (Aperio ImageScope V12.3.3). The slides were also independently assessed in a blinded manner by two board-certified pathologists.

Paraffin-embedded spheroids were sectioned into 4 µm slices, deparaffinized and subjected to antigen retrieval. Spheroids were blocked with 1% bovine serum albumin (BSA) in PBST for 30 min at room temp. Spheroids were stained with primary DCLK1 antibody (as above) overnight at 4°C. The next day, slides were washed with PBS, and spheroid sections were incubated at room temperature in the dark with FITC tagged secondary antibody (Jackson Immuno) at 1:100 dilution for 1 hour. Slides were then rinsed with PBS and counterstained with DAPI and H&E.

### Proliferation and colony formation assay

Cells (5000 cells/well) were plated in 96-well plates, allowed to grow for 24 hours in complete DMEM media containing 10% FBS, and treated with increasing doses of respective vehicle control (DMSO) or DiFiD. Cell viability was measured by enzymatic hexosaminidase assay as described previously (22). Additionally, for colony formation studies, six-well dishes were seeded with 300 viable cells and allowed to grow for 24 hours. The cells were then treated in triplicate with increasing doses of DiFiD. Media was replaced after 24 or 48 hours, removing drug exposure. The cells were incubated an additional 14 days in DMEM medium containing 10% FBS and 2% antibiotic-antimycotic solution (Mediatech Inc, Herndon, VA). Media was changed as needed. The colonies obtained were washed with PBS and fixed using ice-cold methanol for 10 min at room temperature and then washed with PBS followed by staining with Crystal Violet (1% CV in 10% ethanol). The colonies were counted and compared with untreated cells.

### shRNA Transduction

shRNA targeting DCLK1 (NM_004734.2-2100s21c1; sh3 and NM_004734.3-2287s21c1; sh4) and a scrambled shRNA was obtained from Sigma Aldrich (Milwaukee, USA). Lentiviral particles were generated using the pLVX Advanced plasmid system (CloneTech Laboratories Inc, Mountain View, CA) in Lenti-X cells. FaDu cells were transduced with DCLK1 or scrambled shRNA lentivirus, following which clones were grown in DMEM containing 5 µg/mL Puromycin.

### Spheroid formation assay

Single cell suspensions in ultralow attachment plates (Corning, Lowell, MA) of HNSCC (HN5 and FaDu) cells (500 cells/well) were generated. Serum-free growth medium supplemented with EGF (20ng/mL), FGF (20 ng/mL), B27(10 mL in 500 ml of 50X), heparin salt (4 ug/ml) and pen/strep (1% v/v) (Invitrogen) was used to culture the spheroids. Treated samples additionally contained either the vehicle control or IC_50_ concentrations of DiFiD as determined by hexosaminidase assay. Spheroids were centrifuged and 1 mL of ThinPrep PresevCyt Solution (Marlborough, MA) was added to spheroid pellets overnight at 4°C. Spheroids were then paraffin-embedded and processed using immunohistochemistry.

### The Cancer Genome Atlas data analysis

The Cancer Genome Atlas (TCGA) head and neck squamous cancer (GDC HNSC) RNA sequencing (RNA-seq) data were downloaded using the UCSC Xena browser (http://xena.ucsc.edu). HNSC datasets were curated to remove normal adjacent mucosa samples and duplicate samples from metastatic tumors. The data is represented as fragments per kilobase of transcript per million (FPKM) mapped reads, with 271 patients in the low group and 271 in the high group. This was matched to clinical survivorship data from TCGA HNSC phenotype data downloaded from UCSC Xena.

### Molecular docking

All docking calculations were carried out with AutoDock Vina software (23) (Molecular Graphics Lab, Scripps Research Institute, http://vina.scripps.edu/) to analyze DiFiD interactions with the 3D structure of kinase domains of DCLK1 (PDB ID: 5JZN). Default parameters on Autodock tools were used to analyze the docking. Total Kollman and Gasteiger charges were added to the protein and the ligand prior to docking. We used Lamarckian GA to find the best conformations and chose approximately 10 conformations for further analyses. The most stable compound conformation was selected based on the scoring function and the lowest binding energy, and visualized using Pymol (https://pymol.org/2/) (24).

### Cellular thermal shift assay (CETSA)

The ability of DiFiD to interact with and stabilize DCLK1 in cells was determined using CETSA (25). Briefly, cells (8 × 10^6^) were treated with media containing DMSO or DiFiD (5 μM) for 4 hours. After treatment, the cells were aliquoted into PCR tubes and exposed to a temperature gradient. Subsequently, cells were lysed using three repeated freeze-thaw cycles in liquid nitrogen followed by centrifugation. The resultant lysates were then utilized in downstream western blot analyses.

### Animal studies

All protocols were approved by the Institutional Animal Care and Use Committee at the University of Kansas Medical Center. To assess DCLK1 expression at various stages of disease progression, 4-*nitro*-quinoline-oxide (4-NQO,100 ppm in sterile drinking water *ad libitum*) was administered for 16 weeks to C3H mice (n=20) using a previously reported protocol (26). Mice were then given sterile drinking water for 3 weeks, at which time animals were sacrificed and tongues excised. Immunohistochemistry was then used to assess DCLK1 expression.

Five-week-old female Foxn1/nude mice (Charles River Laboratory) were injected with 1×10^6^ HN5 or FaDu cells in the flank. One week following implantation, mice were randomized into two groups with 7 mice per group in the HN5 injected mice and 10 mice per group in the FaDu treated mice. Animals were treated with either vehicle control (2.5% DMSO in water) or DiFiD (2 mg/kg body weight), administered intraperitoneally daily for 15 days. Tumor growth was measured every 2-3 days by a blinded observer measuring tumor diameters using vernier calipers and volume was calculated (Tumor volume = longest dimension × shortest dimension^2^ x 0.52) as previously described (27). At the end of treatment, animals were euthanized, and the tumors were collected, weighed, and processed for downstream analytical assays.

Additionally, we established a patient-derived xenograft from an anal SCC that metastasized to the liver. The tumor was passaged twice through NOD SCID gamma (NSG) mice. ASCC tumors from the third passage were implanted subcutaneously into flanks of 8-week-old mice. Mice were randomized based on the tumor volume into two groups with 10 mice (5 female and 5 male) in the control group and 14 mice (9 female and 5 male) in the DiFiD treatment group. Average tumor volumes across both groups were 50 ± 8.9 mm^3^. Mice were treated daily with either vehicle control (DMSO in water) or DiFiD (2 mg/kg body weight) intraperitoneally daily for 15 days. Tumor growth was measured, and the volume calculated as mentioned before. Fractional tumor volumes were calculated as per previously published protocols (28).

### Statistical analysis

Data are reported as mean ± SEM. Parametric, one-tailed t-test with Welch correction was used to assess significance in all experiments unless stated. Outliers were detected using Graphpad software. Significant outliers were identified using the Grubb’s Test with Alpha=0.05. Significant outliers were removed from further statistical analysis. For *in vivo* studies, repeated measures ANOVA test was employed to assess the level of significance in tumor volumes between treatment arms. For TCGA survivorship comparison, log-rank (Mantel-Cox) test assessed differences between curves. To generate a best expression cut off, patients were stratified into two groups and association between survival and RPKM was examined. The RPKM value that yields the maximum difference between survival of the two groups at the lowest log-rank P-value determined best expression cut off. From this, a high/low expression cut off (0.37) was applied. All statistical calculations were performed on GraphPad Prism software (version 6.03), with significance determined by p<0.05.

## Results

### DCLK1 is upregulated in Squamous Cell Cancers

DCLK1 is associated with pro-survival signaling in various cancers, including colorectal and pancreatic cancers (8, 18, 19). Its expression is elevated in HNSCC tumor samples compared to normal oral epithelial tissue (Fig. 1A and B, p<0.01). TCGA analysis show that increased DCLK1 expression correlated with increased tumor histological grade (Supplementary Fig. S1A), and increase expression was higher in HPV-negative patients (Supplementary Fig. S1B).

**Figure 1.**
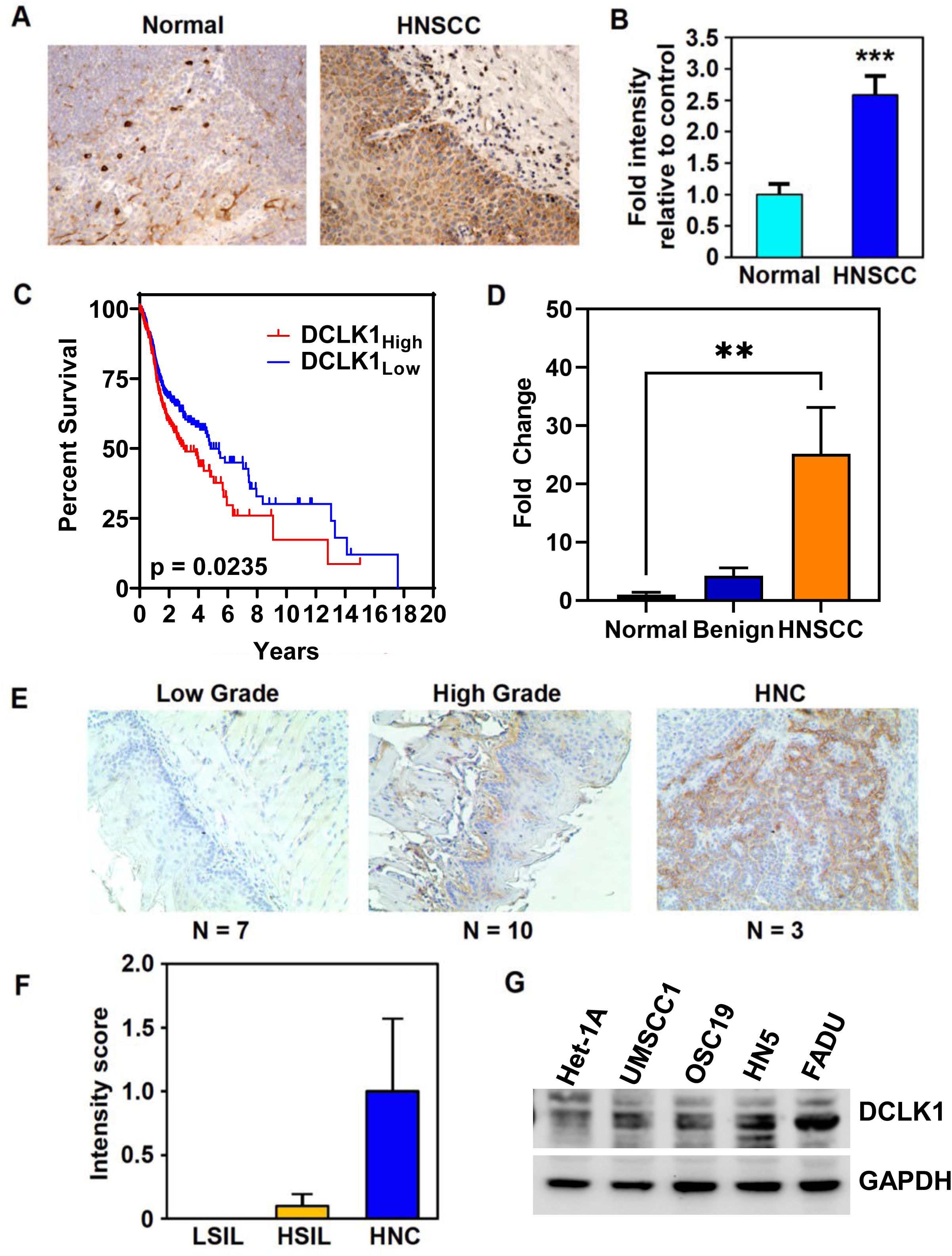
DCLK1 is overexpressed in HNSCC. (A) IHC staining for DCLK1 (arrows) in normal tonsil and HNSCC tissue (B) Fold change in mean intensity of DCLK1 in tissue microarray from HNSCC (n=40) relative to normal oral mucosa (tonsillar) (n=8) samples determined by Leica Aperio ScanScope XT slide scanner (20 images scored per sample) (p<0.003 by two-tailed T- test). (C) TCGA head and neck squamous cancer (HNSC) data demonstrate that high DCLK1 levels trend towards a decrease in overall survival (High, n=192; low, n= 307 *p<0.05). (D) DCLK1 mRNA expression from a cDNA array of HNSCC tumor samples represented as fold change normalized to normal oral mucosa (Normal, n=4; benign, n=3; HNSCC, n=13). (E) Bright field images and (F) scoring of DCLK1 stained oral mucosa tissue, representing low (LSIL, n=7) and high grade (HSIL, n=10), and HNC (n=3) from a 4NQO-carcinogen induced mouse model. (G) Western blot for DCLK1 protein expression in HNSCC cell lines and immortalized non- cancerous squamous esophageal cell line, Het-1A. β-actin used as a loading control. *p<0.05, ***p<0.001.

Kaplan-Meier survival curves generated from TCGA HNSCC mRNA expression datasets demonstrated a trend towards lower overall survival in patients with DCLK1 high expressing tumors (n=192) compared to DCLK1 low expressing tumors (n=307) (Fig. 1C). Patients with DCLK1 low expressing tumors survived on average 4.827 years post diagnosis; patients with DCLK1 high expressing tumors had a shorter median survival of 2.995 years.

To assess correlation of DCLK1 with disease progression, we evaluated the expression of DCLK1 in a cDNA panel, from normal oral mucosa (n=8), benign (n=10), and HNSCC (n=16) patient tissues. DCLK1 was significantly upregulated in cancerous tissue compared to normal (Fig. 1D, p<0.01). To evaluate DCLK1 expression through tumor development, tongue tissue from 4NQO treated mice was collected (n=20). Mice developed various stages of disease with 7/20 mice developing low grade squamous intraepithelial lesions (LSIL), 10/20 developing high grade squamous intraepithelial lesions (HGSIL) and 3/20 developing invasive HNSCC. Higher levels of DCLK1 expression were noted in dysplastic and invasive lesions, with highest expression observed in HNSCC (Fig. 1E and F). We then evaluated DCLK1 expression in HNSCC cell lines; OSC19, HN5, UM-SCC-1, and FaDu and Het-1A, a non-cancerous immortalized esophageal epithelial line. DCLK1 was differentially expressed with the highest expression observed in HN5 and FaDu cells, and lowest expression in Het-1A cells (Fig. 1G). Immunofluorescence revealed DCLK1 to be expressed in HN5 spheroids (Supplementary Fig. S1C). Therefore, HN5 and FaDu were used for subsequent studies.

These data suggest that DCLK1 is elevated in HNSCC compared to normal oral mucosa and is associated with tumor progression. Further, anal squamous cell carcinoma (ASCC) patient samples (n=17) were stained for DCLK1 expression. DCLK1 was more significantly expressed in ASCC tumor compared to normal anal mucosa (Supplementary Fig.S1D and E). We observed increased nuclear staining along the tumor invasive front (Supplementary Fig. S1D).

### DCLK1 suppression inhibits HNSCC growth

To evaluate the antitumor efficacy of targeting DCLK1 in HNSCC, we attenuated DCLK1 levels using two shRNA constructs, sh3 and sh4. DCLK1 mRNA and protein were significantly reduced (Fig. 2A and Fig. 2B) in FaDu cells stably transduced with sh3 or sh4 compared to scrambled controls. DCLK1 has been shown to play a critical role in anchorage independent growth *in vitro*. Hence, we assessed the effects of DCLK1 knockdown on spheroid formation. DCLK1 knockdown suppressed spheroid formation in FaDu cells compared to scrambled controls (p<0.01, Fig. 2C). DCLK1 knockdown also resulted in stunted colony formation (Fig. 2D). Functionally, DCLK1 shRNA knockdown reduced HNSCC migration and invasion *in vitro* (Fig. 2F and Fig. 2G, respectively). To demonstrate that DCLK1 is an important driver of cancer *in vivo*, FaDu cells stably transduced with shRNAs targeting DCLK1 (sh3 and sh4), or a scrambled control shRNA were subcutaneously injected into NOD SCID gamma (NSG) mice. Suppression of DCLK1 resulted in slower growing tumors compared to scrambled controls (Fig. 2I). Tumor lysates were then analyzed by immunoblot for DCLK1, cyclin D1, and Bax (Fig. 2H and Supplementary Fig. 2). Expectedly, DCLK1 expression was diminished in both shRNA groups. Cell cycle regulator, cyclin D1 expression was significantly reduced, while pro-apoptotic, Bax expression was greatly increased in DCLK1 knockdown tumors. Collectively, these data indicate DCLK1 may help drive cancer progression, and DCLK1 suppression may lead to cell cycle arrest and apoptosis.

**Figure 2.**
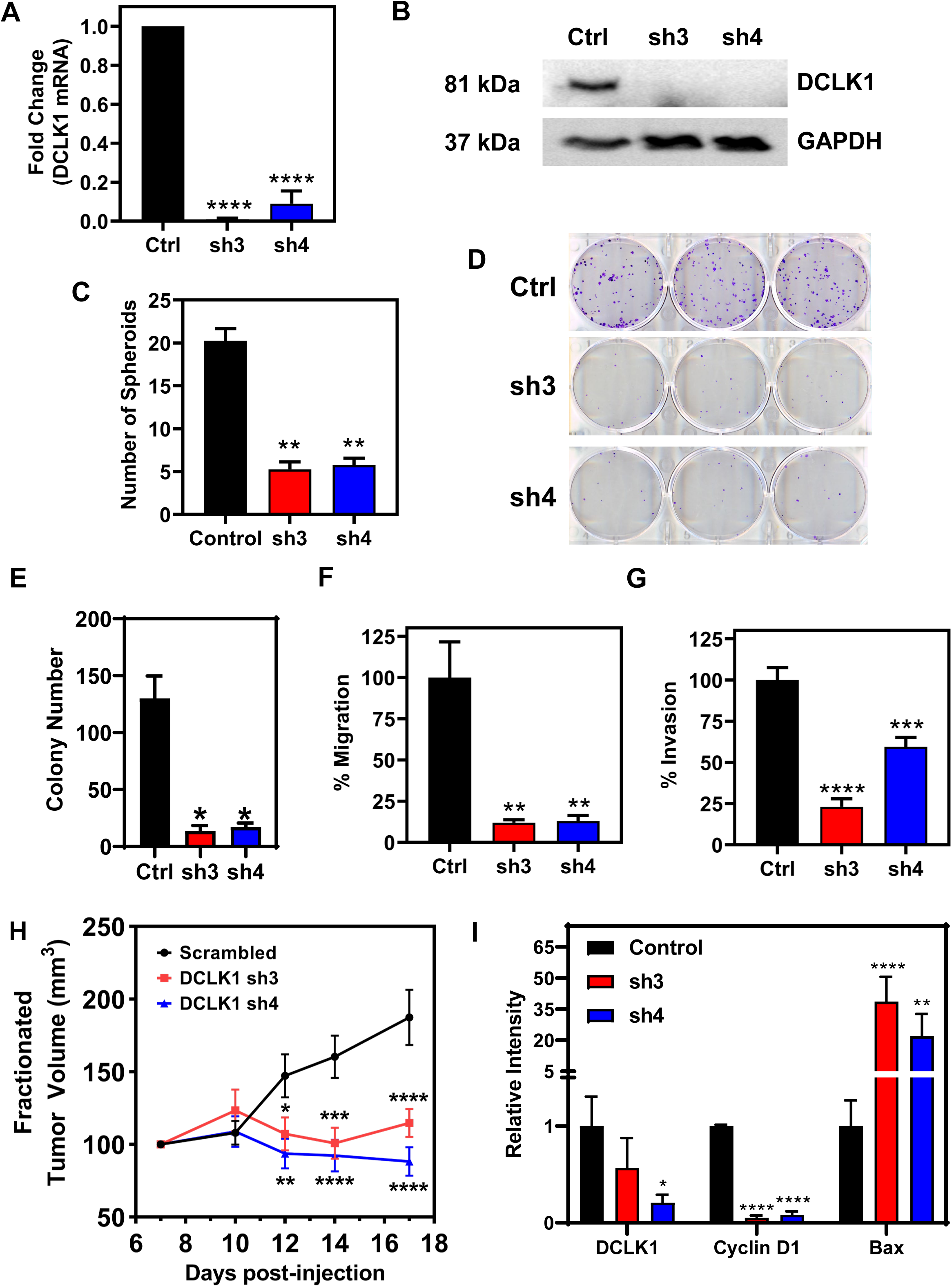
DCLK1 suppression inhibits tumor growth, invasion, and migration. (A) Fold change in mRNA expression of DCLK1 in FaDu cells stably transduced with shRNA targeting DCLK1 (termed: sh3 and sh4) relative to cells transduced with a scrambled control shRNA. Data represent cumulative results from three independent experiments with ± SEM. (B) Western blot of DCLK1 protein in control and stably knocked down DCLK1 sh3 and sh4 FaDu clones. GAPDH served as a loading control. (C) Quantification of spheroid number from spheroid formation assay. Data represent cumulative results from two independent experiments with ± SEM (D) Representative images of colony formation assays for DCLK1 control, sh3 and sh4 FaDu clones. (E) Quantification of colony number. Data represents cumulative results from three independent experiments with ± SEM (F) Migration and (G) invasion assays are normalized for differences in proliferation rates over the duration of the assay. Data represents cumulative results from three independent experiments with ± SEM. (H) Tumor volumes from NSG mice that were inoculated subcutaneously with 1x10^6^ FaDu cells stably transduced with shRNA targeting DCLK1 (sh3 and sh4) or scrambled control shRNA into their right flank (n=10 animals per group). Tumor measurements began 7 days post-injection of cells. Error bars represent ± SEM (I) Quantification of western blot analysis from lysates of a cohort of xenograft tumors. Error bars represent ± SEM (Western blots shown in Supplementary Figure 5A) *p<0.05, **p<0.01, ***p<0.001, ****p<0.0001.

### DiFiD binds to DCLK1 and inhibits its activity

The small molecule, 3,5-bis (2,4-difluorobenzylidene)-4-piperidone (DiFiD) was previously shown to suppress the growth of pancreatic cancer cells *in vitro* and *in vivo* (14). However, the precise target for DiFiD remained unknown. DCLK1 is implicated in pancreatic and colorectal cancer progression. Therefore, we first determined compound-protein interaction. Molecular docking predicted a DiFiD interacts with DCLK1 by forming hydrogen bonds with aspartic acid 533, with a binding energy of -7.9 kcal/mol, as shown in ribbon and surface views (Fig. 3C).

**Figure 3.**
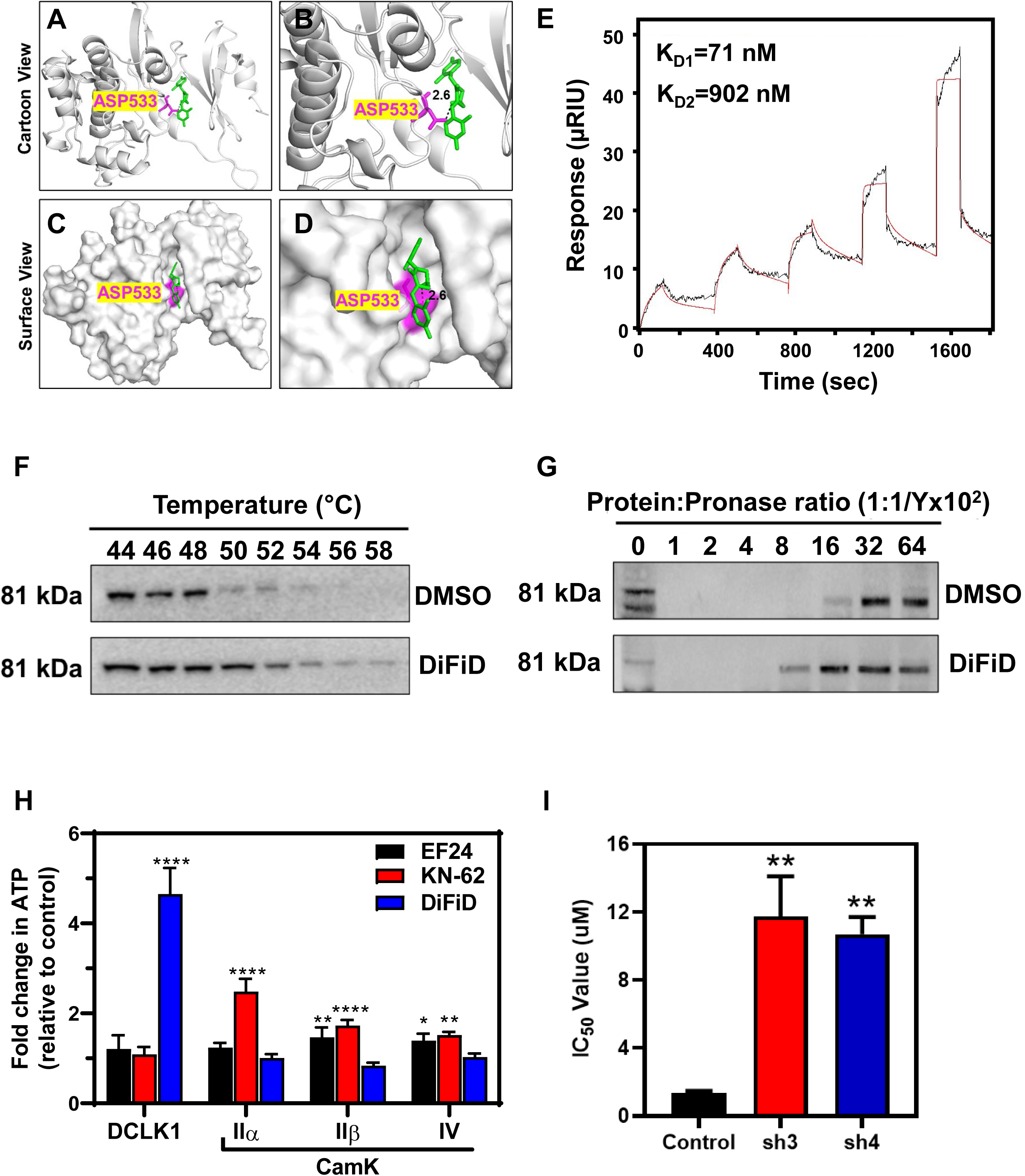
DiFiD binds to DCLK1. Molecular docking studies demonstrating (A) ribbon, (B) magnified ribbon, (C) surface, and (D) magnified surface views of the DCLK1 kinase domain binding to DiFiD with a binding energy (B.E.) of -7.8 Kcal/mol. (E) Surface Plasmon Resonance assay demonstrating the binding affinity of DiFiD analyte to DCLK1 ligand (K_d1_=71 nm, K_d2_ = 902nm). (F) Representative immunoblot of CETSA binding assay of HNSCC (FaDu) treated with 5 μM DiFiD or DMSO for 4 hours then subjected to thermal denaturation. (G) Representative immunoblot of DARTS binding assay of HNSCC (FaDu) treated with 5 µM DiFiD or DMSO, then subjected to enzymatic degradation at various target: enzyme ratios for 15 minutes. (H) *In vitro* kinase assay demonstrating that DiFiD has high selectivity for and greater inhibition of its target DCLK1 relative to other Cam kinase targets. Other inhibitors, EF24 (IKK inhibitor) and KN-62 (pan CamKII inhibitor) demonstrate no significant inhibition of DCLK1. Data represent cumulative results from three independent experiments with ± SEM. **p<0.05, **p<0.01, ***p<0.001, ****p<0.0001 (I) IC_50_ of DiFiD when applied to FaDu cells stably transduced with scrambled control shRNA (control), or shRNA directed against DCLK1-3 (sh3) or DCLK1-4 (sh4). Data represents cumulative data from 3 independent experiments with ± SEM. **p<0.01

Additionally, the binding kinetics of DiFiD and DCLK1 were studied via surface plasmon resonance (SPR). We observed dissociation constants of K_D1_ = 71 nM and K_D2_ =902 nM (Fig. 3E). Taken together this data shows that DiFiD binds DCLK1 with high affinity. To confirm that DCLK1 is a binding target of DiFiD in the cells, we performed cellular thermal shift assay (CETSA) to assess protein stability following thermal denaturation. FaDu cells were treated with 5 µM DiFiD at 37°C for 4 hours. Cells were aliquoted into equal volumes, subjected to a thermal gradient and DCLK1 expression was then evaluated by western blot. Thermal denaturation of DCLK1 occurred at 54°C in the DMSO treated control group, which increased to 58°C in the presence of DiFiD (Fig. 3F). To further validate DiFiD binding with DCLK1, we performed the drug affinity responsive target stability (DARTS) assay. Briefly, FaDu cell lysates were incubated with DMSO or DiFiD (5 µM) for 30 minutes followed by treatment with increasing concentration of pronase. Starting with 0 mg/mL stock solution of pronase, we incubated cells with increasing protein:pronase ratio. We observed that DiFiD protected DCLK1 from protease-mediated degradation, as DCLK1 expression was extended to 1: 1/800 protein: pronase ratio compared to 1: 1/1600 in the DMSO control arm (Fig. 3G). Altogether, these data suggest that DiFiD binds to DCLK1.

To confirm DiFiD specifically interacts with the kinase domain of DCLK1, we performed CETSA analyses. For this, we stably expressed the N-terminal DCX domain (lacking the kinase domain) in HNSCC cells (Supplementary Fig. S3A and S3B). To detect DCX domain fragments in FaDu cells, we utilized DCLK1 antibody produced using residues near the N-terminus. This specifically recognizes DCLK1 protein isoforms containing 82 kda DCX sequences. The DCX fragment we developed had a predicted molecular weight of 51 kDa. We found that cells expressing the 51 kDa N-terminal DCX domain did not exhibit thermal stabilization when treated with DiFiD (Supplementary Fig. S3C and S3D). This suggests that DiFiD does not stabilize DCLK1 lacking the kinase domain. Therefore, we conclude that DiFiD preferentially binds to the kinase domain of DCLK1, further confirming molecular docking predictions.

DCLK1 belongs to the family of kinases with homology to calmodulin kinases, but it does not depend on Ca2+/calmodulin for its kinase activity (29). Since DiFiD interacts with the kinase domain, we next assessed the effect of DiFiD on DCLK1 kinase activity by performing an *in vitro* kinase assay. Here, we incubated recombinant DCLK1 with GS peptide as substrate to assess the ability of DCLK1 to phosphorylate the peptide (18, 19). Along with DiFiD, we also tested the ability of EF24 (IKK inhibitor) and KN62, a pan-CaMKII inhibitor. We observed greater inhibition of DCLK1-mediated ATP consumption following incubation with DiFiD, than with either EF24 or KN-62 (Fig. 3H). To determine the specificity of DiFiD to DCLK1, we also performed studies with CaMKIIα, CaMKIIβ, and CaMKIV. We observed that DiFiD did not affect the activities of these CaMK proteins. These data demonstrate a higher selectivity and specificity of DiFiD for DCLK1.

Since DiFiD has favorable binding to DCLK1, inhibits its kinase activity, and is relatively ineffective at reducing Het1A proliferation (Supplementary Fig. 4B), we hypothesized that cells expressing lower levels of DCLK1 would be resistant to DiFiD. Consequently, we performed hexosaminidase assays with cells where DCLK1 was knocked down using specific shRNA. There was an increase in their respective half maximal inhibitory concentration (IC_50_) values from 1.3 μM to greater than 11 μM compared to control cells (Fig. 3I). These data demonstrate DiFiD activity is dependent upon the presence of DCLK1, and DiFiD has poor activity against cells with low DCLK1 expression. These data further suggest that DCLK1 is a highly specific direct target of DiFiD.

### DiFiD demonstrates potent cytotoxicity in HNSCC *in vitro*

Since DiFiD inhibits DCLK1 activity, we sought to determine the effect of DiFiD on HNSCC cancer cell viability. We observed that DiFiD inhibits HNSCC viability in a dose and time-dependent manner (Fig. 4A and Supplementary Fig. S3A). We identified an effective dose to assess the mechanism of action. The IC_50_ was measured by hexosaminidase assay and observed within 48 hours at concentrations of 750 nM and 1.5 µM in HN5 and FaDu cell lines, respectively. In addition, we tested toxicity of DiFiD on an immortalized non-cancerous cell line, Het1A (30) (Fig. 1G). We observed that the IC_50_ for Het1A at 48 hours was 9 µM, a 6-12-fold increase compared to FaDu and HN5 cells, respectively (Supplementary Fig. S4B). In addition, we observed that DiFiD attenuates spheroid growth in both HN5 and FaDu cell lines when treated at IC_50_ concentrations, suggesting that it affects anchorage independent growth (Fig. 4B). We next determined the effect of DiFiD on clonogenicity of HNSCC cells. Initial studies with IC_50_ doses demonstrated complete suppression of colony formation. However, when treated at lower doses of 93.75 nM (1/8th IC50) and 187.5 nM (1/4th IC50) for 48 hours, we observed a dose dependent decrease in both the number and size of colonies for HN5 cells (Fig. 4C and D, p<0.01). Similarly, FaDu cells treated with a 1/4th (375 nM) and 1/8th (187.5 nM) IC_50_ dose of DiFiD exhibited a significant inhibition in colony number and size (Fig. 4E and F, p<0.01). These data suggest that DiFiD has potent cytotoxic effects on HNSCC that are long lasting.

**Figure 4.**
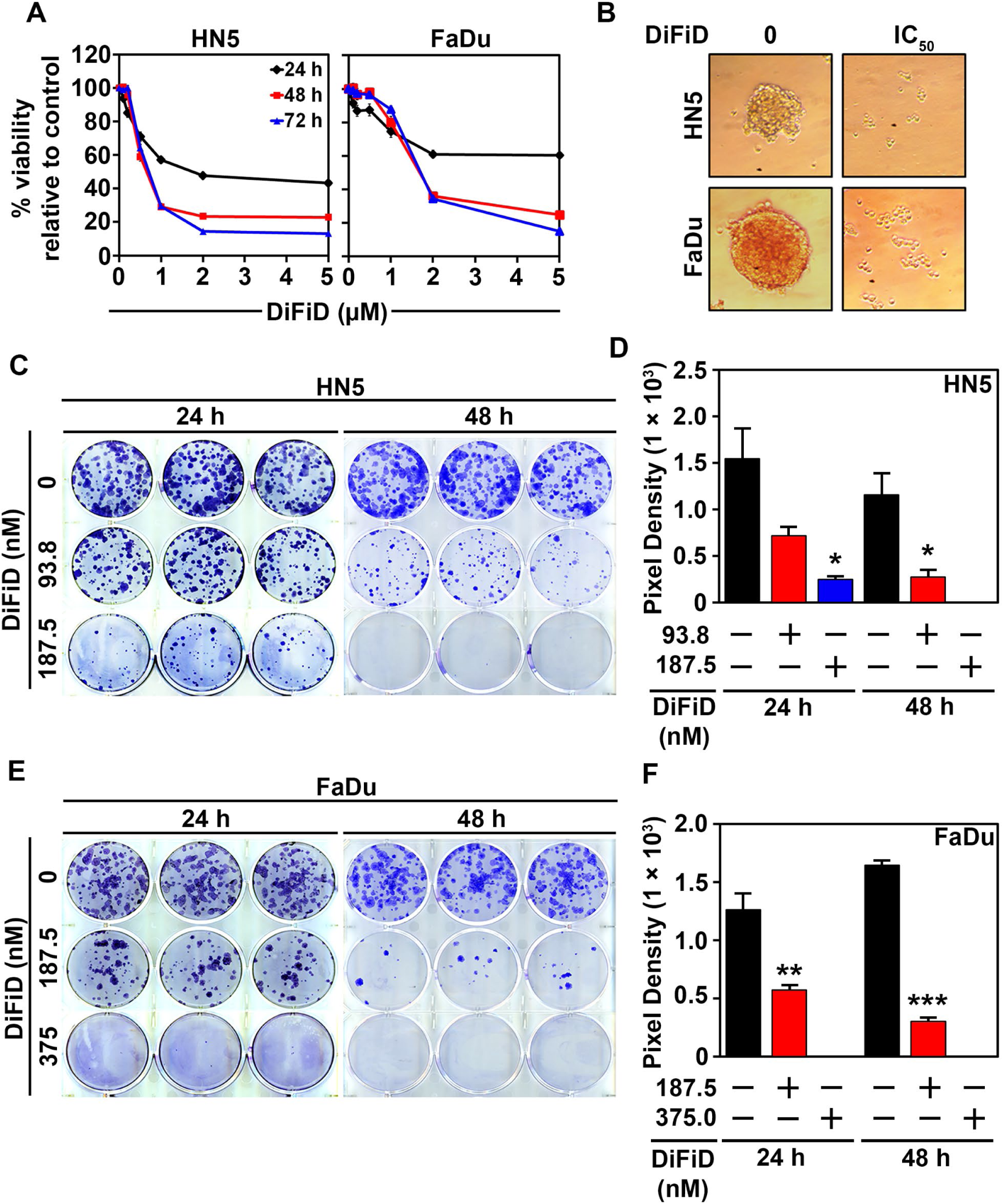
DiFiD inhibits HNSCC proliferation in 2-D and 3-D culture models. (A) Hexosaminidase viability assay of HNSCC (HN5 and FaDu) treated for 24, 48, and 72 hours at increasing concentrations of DiFiD (0, 0.125, 0.25, 0.5, 1, 2, 5 µM). Figure is representative of three experimental replicates with ± SEM. (B) DiFiD abrogated spheroid growth in HNSCC (HN5 and FaDu) at IC_50_ concentrations. (C-F) Colony formation assay was performed to assess long term efficacy of 24 h or 48 h DiFiD treatment. Colony formation assay and quantification of pixel density representing the colony size using ImageJ of HN5 (C, D) and FaDu (E, F), 14 days post DiFiD treatment. Graphs represents cumulative results from three independent experiments with ± SEM. *p<0.05, **p<0.01, ***p<0.001.

### DiFiD induces G2/M arrest and apoptosis

To further characterize the effects of DiFiD treatment on HNSCC proliferation, we used flow cytometry to evaluate cell cycle progression. HN5 and FaDu cell lines were treated with DiFiD. Within 24 hours, DiFiD treatment increased the percentage of cells arrested at the G_2_/M phase (Fig. 5A and Supplementary Fig. S5). At 48 hours, there was an increase in the sub G_0_, or fragmented DNA stage following treatment, suggesting increased cell death. To confirm G_2_/M arrest, western blot analysis was performed for G_2_/M associated proteins cyclin B1 and cell division cycle protein 2 (CDC2). HN5 and FaDu cancer cells were treated with IC_50_ doses of DiFiD for 24 hours, in which time dependent increases in cyclin B1 and simultaneous decreases in CDC2 expression were observed in the treatment arm (Fig. 5B and 5C). We also performed western blot analysis for pro- and cleaved-PARP protein. DiFiD treatment induced PARP cleavage in both HN5 and FaDu cell lines (Fig. 5B and 5C). This suggests that DiFiD further induces apoptosis in HNSCC cancer cell lines associated with a G_2_/M arrest. This was confirmed by Annexin V/PI staining that demonstrated an increased percentage of DiFiD treated cells were in the apoptotic and dead fractions compared to vehicle control (Fig. 5D). To assess the role of caspases in the induction of apoptosis by DiFiD, we performed caspase 3/7 assay to assess effector caspase activity. DiFiD induced a significant upregulation in the activity of the effector caspases (Fig. 5E, p<0.001). Taken together, these data confirmed that DiFiD mediated suppression of DCLK1 activity lead to mitotic catastrophe of HNSCC.

**Figure 5.**
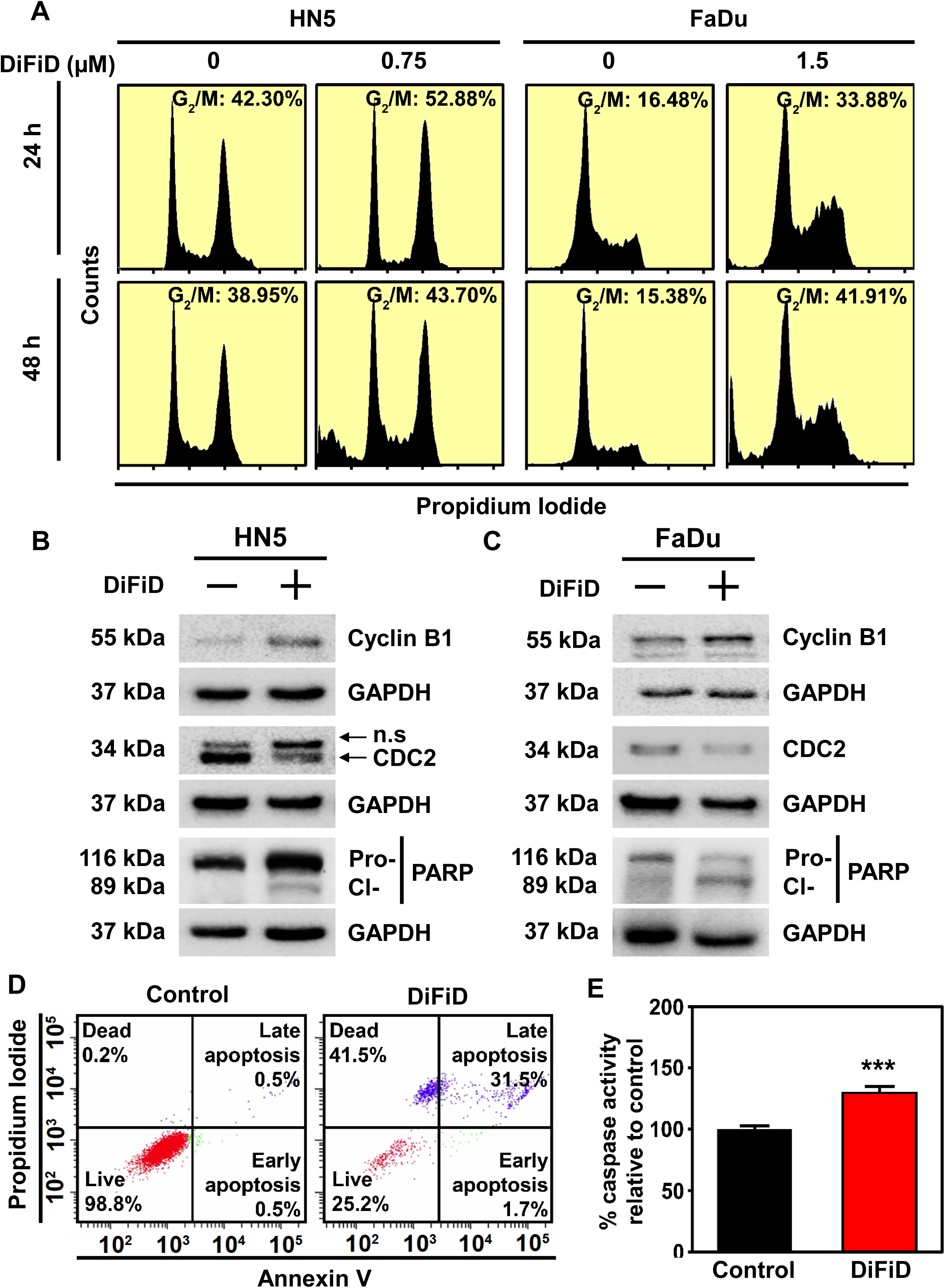
DiFiD induces G_2_/M arrest and apoptosis. (A) Cell cycle analysis was performed in HN5 and FaDu cell lines after 24 or 48 h treatment with IC_50_ concentrations of DiFiD, (HN5=750 nM; FaDu=1.5 µM). Figure is representative of three independent studies. (B) Representative immunoblot of HN5 cells treated with 750 nM DiFiD for 24 hours demonstrates expression of cyclin B1, CDC2, Pro- and cleaved-PARP protein with GAPDH used as a loading control. (C) Representative immunoblot of FaDu cells treated with 1.50 µM DiFiD for 24 hours demonstrates expression of cyclin B1, CDC2, Pro- and cleaved-PARP protein with GAPDH used as a loading control. (D) Annexin V assay of HN5 cells following treatment with DiFiD at the IC_50_ dose. HN5 cells were treated with DiFiD at the IC_50_ dose for 48 h, stained with Annexin V (FITC) and PI, and analyzed by flow cytometry. (E) Caspase 3/7 assay demonstrates caspase cleavage in HN5 cells 48 h post treatment with DiFiD. Data represents cumulative results from three independent experiments with ± SEM ***p<0.001.

### DiFiD has antitumor effects *in vivo*

To determine the *in vivo* antitumor activity of DiFiD, we treated FaDu and HN5 subcutaneous tumors in *Foxn1^nu/nu^*mice. Briefly, HN5 or FaDu cells were injected subcutaneously into the flanks of nude mice and were subsequently treated with DiFiD at 2 mg/kg/day for 15 days (Fig. 6A). DiFiD treatment significantly inhibited tumor growth in HN5 xenografts compared to vehicle control (DMSO) treated tumors (n=7, p<0.05, Fig. 6B and Supplementary Fig. 6A). Similarly, with FaDu xenograft subcutaneous tumors, DiFiD treatment significantly reduced tumor growth compared to the vehicle control treatment arm (n=10, p<0.05, Fig, 6C and Supplementary Fig. 6E). Lastly, the antitumor effect of DiFiD was tested in an ASCC patient-derived xenograft model. DiFiD treatment significantly reduced PDX tumor growth as evidenced by tumor volume and weight of ASCC PDX tumors when compared to the control group (p<0.05, Fig. 6D and p<0.001, Fig. 6E). These preclinical data indicate that DiFiD is a promising, well-tolerated, therapeutic agent for the management of SCC.

**Figure 6.**
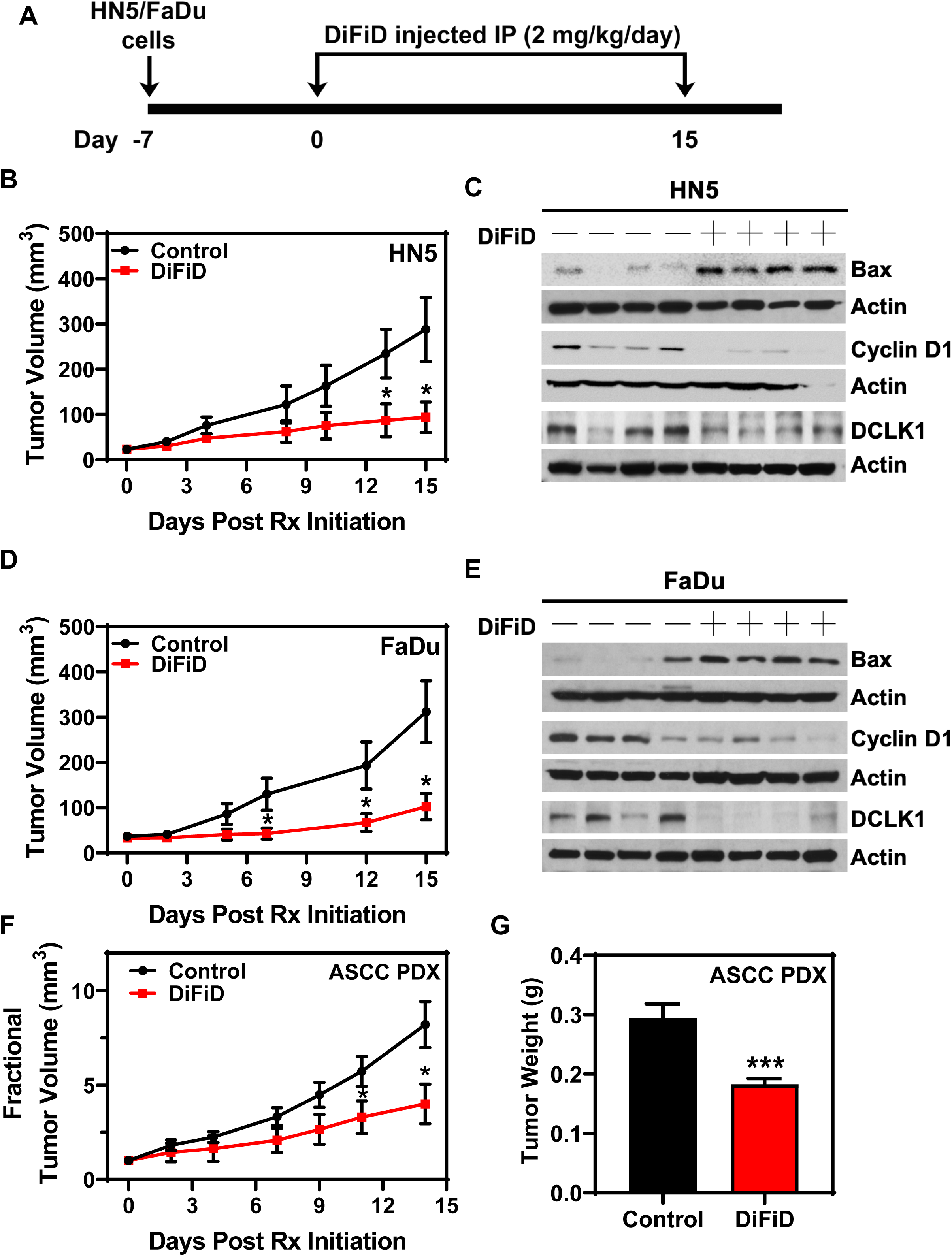
DiFiD inhibits SCC growth *in vivo*. (A) DiFiD treatment protocol for xenograft studies. HN5/FaDu(1 x 10^6^) cells were inoculated subcutaneously into the right flank of athymic *Foxn1^nu/nu^* mice. DiFiD (2 mg/kg) or vehicle control (DMSO) was administered by intraperitoneal injection once daily for 15 days. (B) HN5 tumor volumes (n=7 mice/group) are depicted. (C) HN5 xenograft tumors were subjected to biomarker analyses. Representative immunoblots from a small cohort of HN5 xenograft tumors were probed for Bax, cyclin D1 and DCLK1. (D) FaDu tumor volumes (n=10 mice/group) are depicted. (E) Representative immunoblots from a small cohort of FaDu xenograft tumors were probed as in (C). ASCC PDX was transplanted into the flanks of NSG mice. DiFiD (2 mg/kg) or vehicle control (DMSO) was administered by intraperitoneal injection once daily for 15 days. (F) Fractional tumor volume and weight (G) are depicted (n=10 mice/group for control and n=14 mice/group for DiFiD treatment). Error bars represent ± SEM. *p<0.05, ***p<0.001.

To elucidate the molecular mechanism whereby DiFiD exerts its antitumor effects on HNSCC, we analyzed xenograft tumors using western blotting. Tumor samples were subjected to electrophoresis, and subsequently, expression of Bax, Cyclin D1, and DCLK1 were determined. DiFiD significantly induced the expression of Bax in HN5 (Fig. 6F and Supplementary Fig. 6B) and FaDu (Fig. 6G and Supplementary Fig. 6F) tumor samples relative to vehicle control. Additionally, DiFiD significantly reduced expression of Cyclin D1 (Fig. 6F and Supplementary Fig. 6C) and DCLK1 (Fig. 6F and Supplementary Fig. 6D) in HN5 tumors, suggesting inhibition of proliferation and cell cycle entrance. FaDu tumor xenografts also demonstrated decreased expression of cyclin D1 (Fig. 6G and Supplementary Fig. 6G) and DCLK1 (Fig. 6G and Supplementary Fig. 6H), further substantiating our observations *in vivo*. Taken together, these studies demonstrate that DiFiD inhibits tumor growth by targeting DCLK1 and inducing cellular apoptosis.

## Discussion

SCCs from the head and neck and anorectal regions are highly aggressive malignancies with high rates of local recurrence, distant metastasis, and poor clinical outcomes, including reduced survival. Current standard of care include surgery followed by chemoradiotherapy. However, despite aggressive treatment, the survival rate remains low highlighting a significant unmet medical need in SCC patients. In this article, we demonstrate that DCLK1 is a clinically relevant target for HNSCC and ASCC. DCLK1 is well characterized as a reserve, stress induced stem cell marker in pancreatic and colorectal cancers (5, 6). Using multiple platforms, we demonstrate DCLK1 expression is significantly upregulated in HNSCC compared to normal oral mucosa of patient tissues. Furthermore, high expression of DCLK1 correlates with poor clinical outcomes. These findings agree with a recent report associating high DCLK1 levels with poor HNSCC patient survival (4).

Genetic knockdown of DCLK1 has demonstrated promising findings in neuroblastoma, colorectal, and pancreatic tumors (31, 32). DCLK1 knockdown triggers apoptosis and inhibits proliferation, mitochondrial function, and ATP synthesis in neuroblastoma cells (32). Furthermore, DCLK1 siRNA nanoparticle delivery to colorectal and pancreatic tumor xenografts resulted in the significant inhibition of tumor growth with seemingly high tolerance (31). Our data presented herein supports these previous findings, as we observed decreased spheroid growth, colony formation, migration, and invasion following knockdown of DCLK1. As such, due to the high expression of DCLK1 in HNSCC tumor tissues and cell lines, we postulated that DCLK1 may be a potential therapeutic target for HNSCC.

There are no known DCLK1 inhibitors in development, in fact, few compounds have been reported to inhibit DCLK1, and none with high specificity. Yet, in a single study, Weygant et al., in targeting leucine-rich repeat kinase 2 (LRRK2), with the small molecule inhibitor LRRK2-IN- 1, reported inhibition of DCLK1, and subsequent attenuation of HCT116 (colon) and AsPC-1 (pancreatic) growth, and invasiveness (33). However, direct binding of DCLK1 by LRRK-IN-1 was not examined in this study and the effects are most likely due to the indirect effects of LRRK inhibition on DCLK1 activity (34). Furthermore, several kinase inhibitors with anti-tumor activity, such as XMD8-92 (MAPK7 inhibitor), BI-2536 (PLK1 inhibitor), and TAE-684 (ALK inhibitor), demonstrate non-specific activity, with a comparable affinity towards DCLK1 as much as their target kinases (35). Therefore, while inhibition of DCLK1 may play a role in the therapeutic activity of these compounds, it is highly valuable to identify a specific DCLK1 inhibitor for future clinical applications.

Previously, we reported that DiFiD shows antitumor activity towards pancreatic cancer cells [11]. However, the direct target for DiFiD was not elucidated. We used an *in vitro* kinase assay to demonstrate that DiFiD effectively inhibits DCLK1 kinase activity, but not related CaMK family members, suggesting that the compound is a specific competitive inhibitor of DCLK1. We further confirmed a direct interaction through *in vitro* binding assays and SPR analysis. The CETSA and DARTs binding assays involved the uptake of the compound by cells before thermal or enzymatic denaturation, respectively (25). We showed that DCLK1 was robustly stabilized by DiFiD. Additionally, cells stably transfected with expression plasmids containing only the DCX domain did not exhibit thermal stabilization as was observed with full length DCLK1 containing the kinase domain. Interestingly, we found that DCX domain required higher temperatures to denature compared to full length protein. This is likely due to enhanced microtubule association and stabilization of the domain, as this domain in DCLK1 is critical for its microtubule binding and polymerization activity (7, 36). SPR analysis, which is the gold standard for target interactions, identified strong DiFiD:DCLK1 binding with low nM concentration equilibrium dissociation constants K_D_. Taken together, these data demonstrate that DiFiD has high selectivity for DCLK1 with binding sites located in the c-terminal kinase domain.

We demonstrated that DiFiD induces cell-cycle arrest and apoptosis, attenuating HNSCC proliferation and colony formation *in vitro*, and tumor growth *in vivo*. Het1A, non-cancerous cells expressing low amounts of DCLK1 in comparison to the HNSCC cell lines, HN5 and FaDu, demonstrated lower DiFiD mediated toxicity. When we knocked-down DCLK1 in FaDu cells using shRNA, we observed a 10-fold decrease in DiFiD toxicity, further demonstrating the specificity of DiFiD for DCLK1. DiFiD increased cell cycle arrest in the G_2_/M phase resulting in mitotic catastrophe, data consistent with previously published studies (14, 33). This is further supported by our observation of a significant increase in the sub-G_0_ population following treatment with DiFiD. Moreover, we observed an upregulation of the G_2_/M associated protein, cyclin B1 and a decrease in cdc2, proteins that are abnormally regulated when cells undergo mitotic catastrophe (37, 38). These data corroborate our previous reported results demonstrating the induction of p21 and a reduction in cyclins A1 and D1 following DiFiD treatment in pancreatic cancer (14).

ASCC has an incidence rate of 0.2-4.4 per 100,000 people per year which has risen worldwide over the last three decades, especially in homosexual men (35 per 100,000 per year) and those with HIV (75-135 per 100,000 per year) (39, 40). Despite these rising numbers, the standard of care treatment for this cancer comprised of fluorouracil and mitomycin C has remained essentially unchanged since its inception (41). The addition of intensity-modulated radiation therapy and cisplatin has shown similar efficacy to fluorouracil and mitomycin C in a large, randomized trial (42, 43). The relative rarity of ASCC presents challenges in conducting pivotal clinical trials. In addition, validated preclinical models of ASCC that accurately replicate clinical observations are limited. Existing preclinical models of ASCC include a cell line derived from a lymph node metastasis, two transgenic mouse models and a xenograft from a single patient (44–47). Here we present a patient derived xenograft model and provide preliminary evidence of a promising, new therapeutic target for the management of ASCC.

In our studies, we observed significant antitumor effects in SCC preclinical models following treatment with DiFiD. DCLK1 is expressed in various normal cell types, including neurons, osteoblasts, and colon stem cells (48), and is involved in physiological processes, including retrograde transport, neuronal migration, and neurogenesis. The data presented in this article suggest that DiFiD is well tolerated at doses that demonstrated antitumor activity, as mice maintained normal weight gain and ambulation. DiFiD tolerance was observed in an earlier study (14), further supporting the notion that it is well tolerated at the doses used to inhibit cancer growth.

In conclusion, for the first time we demonstrated that DiFiD binds to DCLK1 and inhibits SCC growth *in vitro* and *in vivo*. Thus, our results strongly establish DCLK1 as a clinically relevant therapeutic target for SCC.

## List of abbreviations

ASCC: Anal Squamous Cell Carcinoma
CETSA: Cellular Thermal Shift Assay
DARTS: Drug affinity responsive target stability assay
DCLK1: Doublecortin-Like Kinase 1
DiFiD: 3,5-bis (2,4-difluorobenzylidene)-4-piperidone
DMSO: Dimethyl Sulfoxide
HGSIL: High grade squamous intraepithelial lesions
HNSCC: Head and Neck Squamous Cell Carcinoma
HPV: Human Papilloma Virus
LRRK2: leucine-rich repeat kinase 2
LSIL: Low grade squamous intraepithelial lesions
NSG: NOD SCID gamma mice
PDX: Patient derived xenograft
RPKM: Reads Per Kilobase of transcript, per Million mapped reads
SCC: Squamous Cell Carcinoma
TCGA: The Cancer Genome Atlas

## Acknowledgements

We would like to thank The University of Kansas Cancer Center Biospecimen Repository Core Facility for their services in providing high quality tissue samples.

## Authors’ contributions

DS curated, analyzed and interpreted data, and was a major contributor in writing the manuscript. LA curated, analyzed, and interpreted data, and was a major contributor in writing the manuscript. PD curated data for *in silico* binding, DARTS and CETSA binding assays, and contributed to manuscript writing and editing. BO curated, and analyzed data regarding DCLK1 knockdown in HNSCC cells and animal models. VS curated, and analyzed spheroid formation and viability assay data, and assisted in writing of the manuscript. SP curated western blots for animal studies. AS assisted in curation, and analysis of TCGA datasets and manuscript writing/editing. DS curated, analyzed and interpreted data for cDNA array. PS curated data for IHC staining, assisted with lentiviral production, and manuscript editing. SC assisted with animal data curation and colony maintenance. JN assisted with animal data curation and colony maintenance. DK curated and analyzed data for DCLK1 kinase activity. PR curated IF staining of HNSCC spheroid cultures. BR assisted with DCLK1 cloning and truncation experiments. MS assisted with curation of data for viability assays and migration and invasion datasets. RA Study consultant. MO IHC scoring SG Statistics consultant. JA Tissue collection. SU Study consultant. SW Study consultant related to drug delivery. OT Secondary IHC scoring consultant SP Oversaw production of DiFiD and drug study consultant. SA Interpreted datasets, major contributor in writing manuscript, oversaw study design. ST interpreted datasets, major contributor in writing manuscript, oversaw study design. All authors read and approved the final manuscript.

## Declarations

### Ethics approval and consent to participate

Human tissue explants were collected after signed consent from patients by the University of Kansas Cancer Center, Biospecimen Repository Core Facility. The Repository complies with all regulations governing the protection of human subjects including CFR 45 Part 46, HIPAA, and the KUMC Human Subject Committee (HSC). Tissue collection and use were carried out under protocol HSC #5929. All animal protocols were approved by the Institutional Animal Care and Use Committee at the University of Kansas Medical Center.

### Availability of data and materials

The datasets generated or analyses during this study are available from the corresponding author on reasonable request.

## Supplementary Figure Legends

**Supplementary Figure 1.**
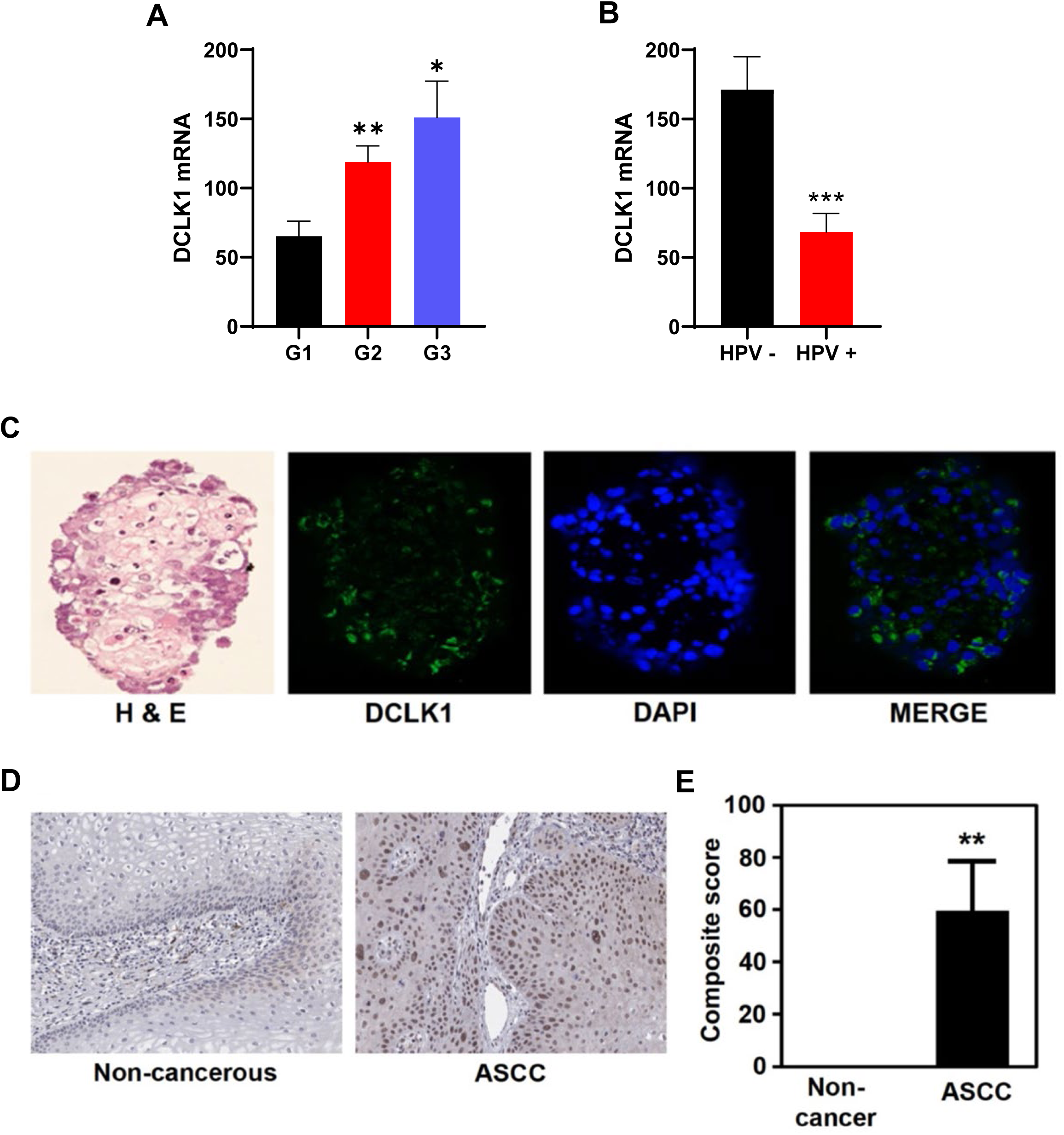
(A) DCLK1 mRNA expression by histological tumor grade represented as fragments per kilobase of transcript per million (FPKM) mapped reads, obtained through TCGA analysis (G1, n=64; G2, n=312; G3, n=125) (B) DCLK1 mRNA expression in HPV negative (n=74) and HPV positive tumors (n=42) represented as fragments per kilobase of transcript per million (FPKM) mapped reads, obtained through TCGA analysis (C) HN5 spheroids stained with H&E, or DCLK1 (green) and DAPI (blue). (D) Representative images of DCLK1 staining in non- cancerous anal mucosa (n=3) and ASCC (n=14). (E) Cumulative results of composite score (intensity x % positivity) of DCLK1 staining in anal mucosa samples. *p<0.05, **p<0.01, ***p<0.001

**Supplementary Figure 2.**
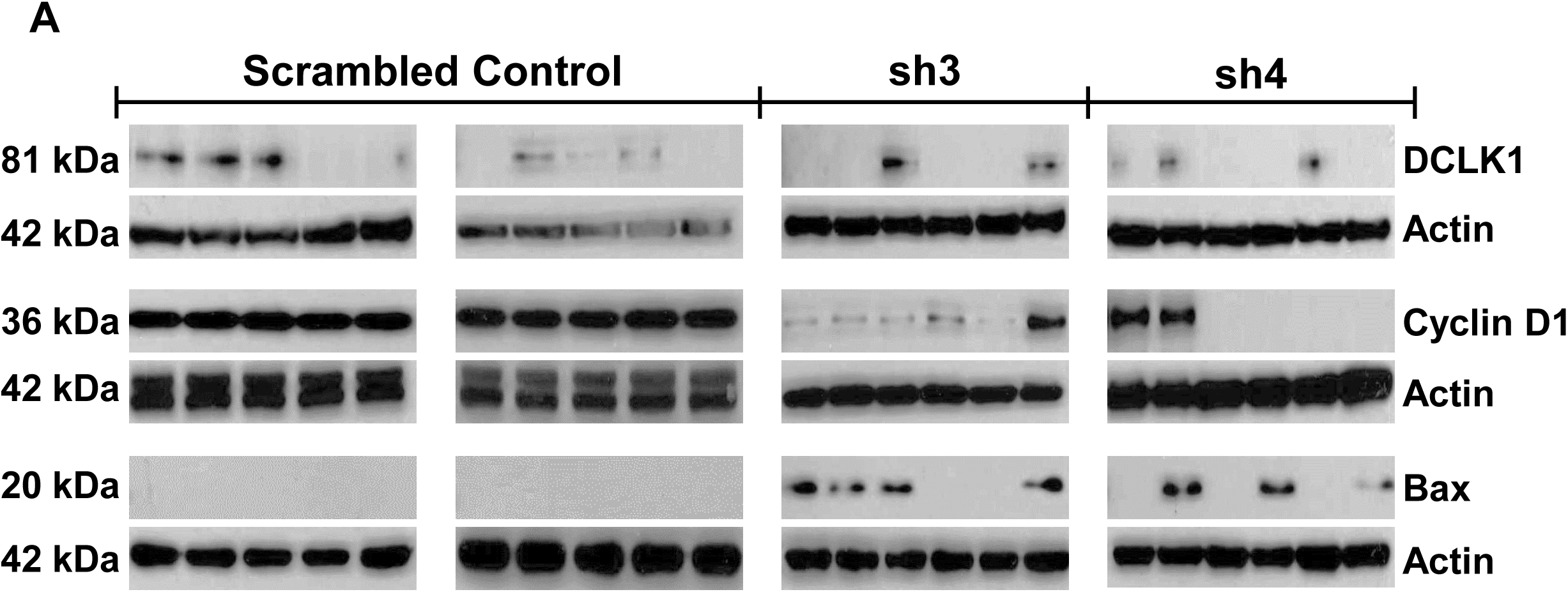
(A) Western blots from xenographic tumor lysates. 1x10^6^ Fadu cells stably transfected with shRNA targeting DCLK1-3 (sh3), DCLK1-4 (sh4) or scrambled control shRNA were injected subcutaneously into NSG mice.

**Supplementary Figure 3.**
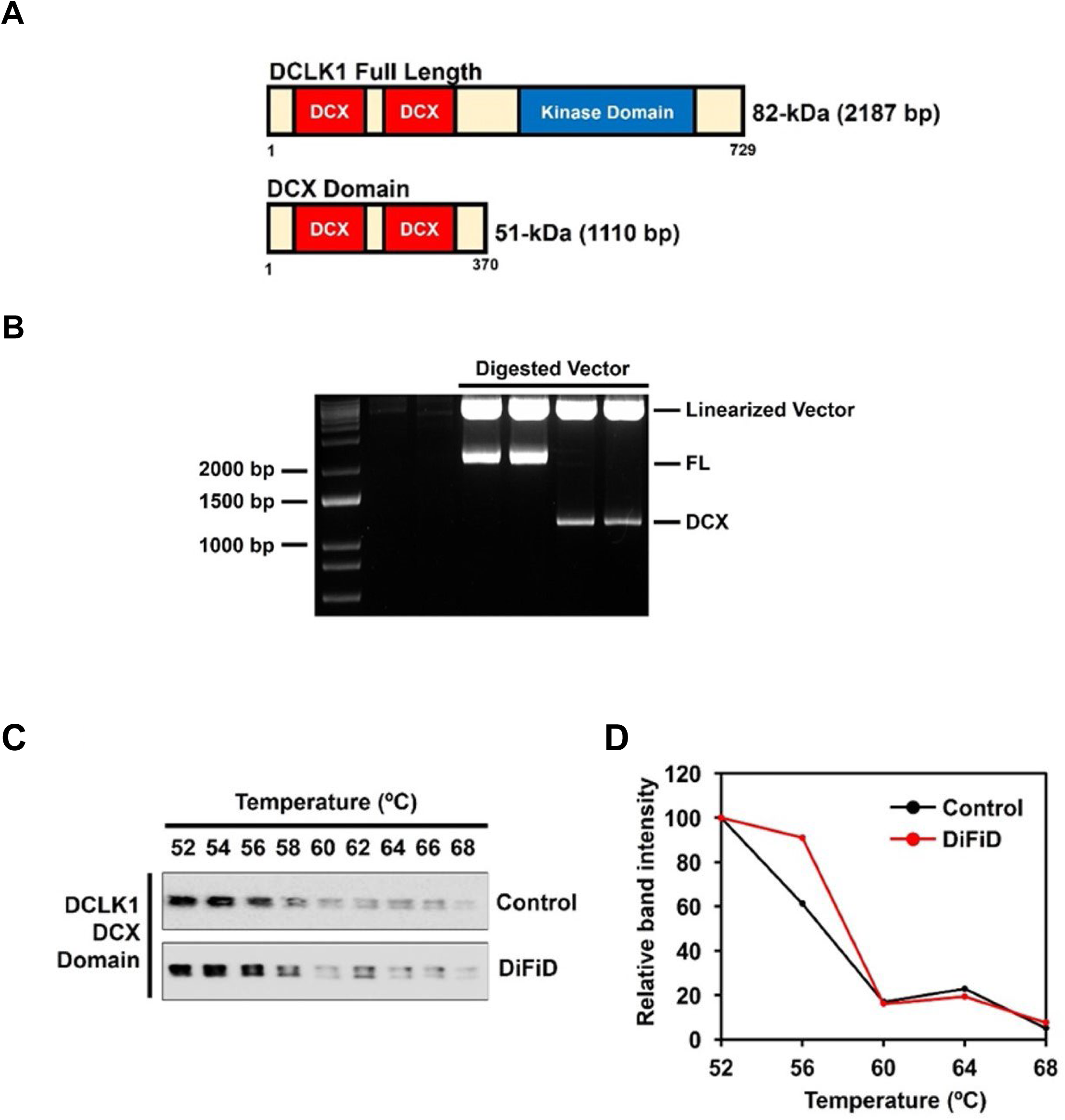
(A) Diagram of DCLK1 full length and DCX domain fragments. (B) Restriction digested plasmid constructs expressing either full length (FL) or DCX domain (DCX) fragments. (C) Western blot analysis of CETSA samples from DCX domain expressing FaDu cells treated with DMSO or DiFiD. (D) Cumulative densitometric evaluation of CETSA samples as described in (C).

**Supplementary Figure 4.**
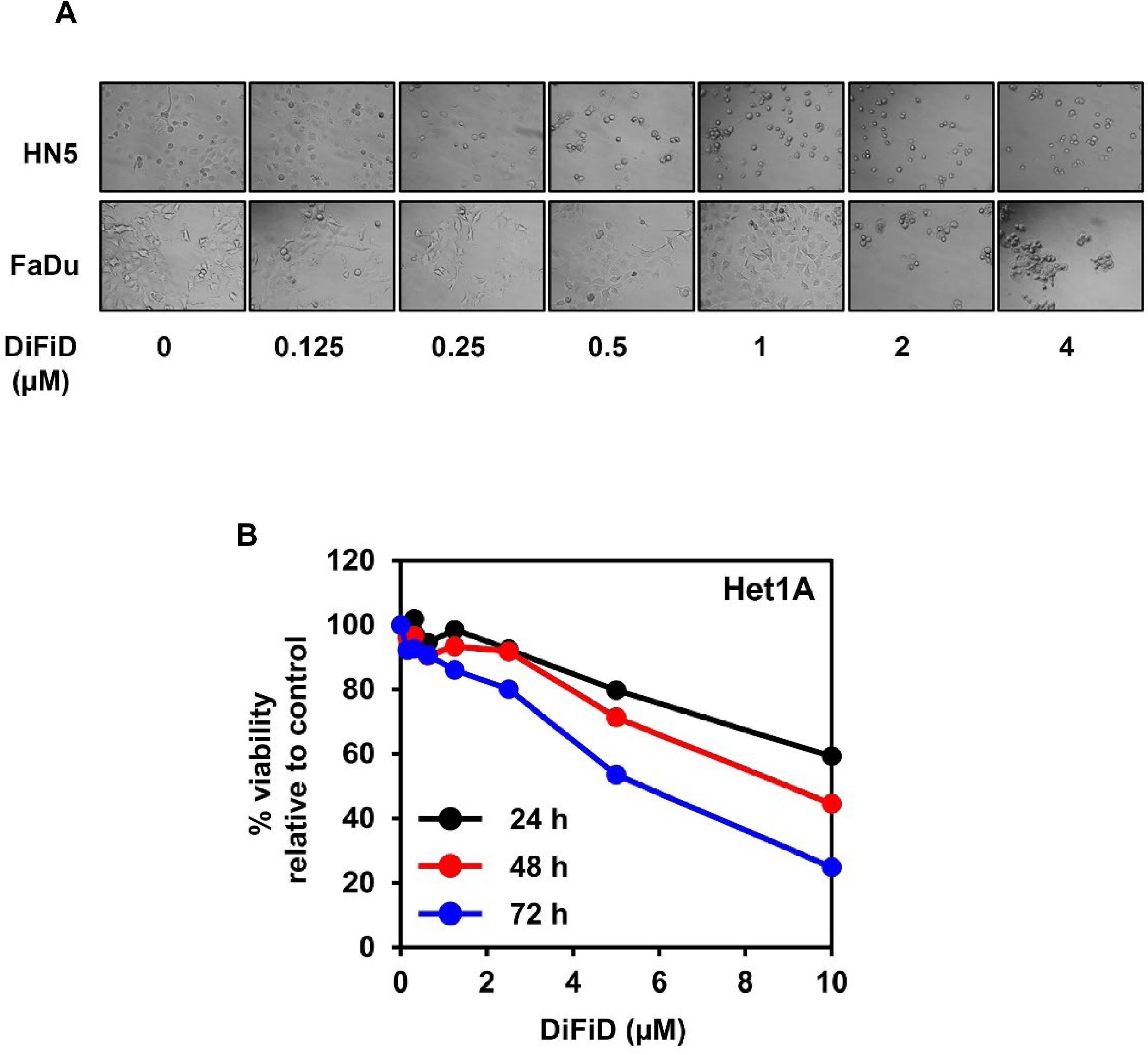
(A) Bright field images of HN5 and FaDu cells at 100X magnification treated with increasing doses of DiFiD for 48 h. (B) Hexosaminidase assay for Het1A cells treated with DiFiD for up to 72 hours.

**Supplementary Figure 5.**
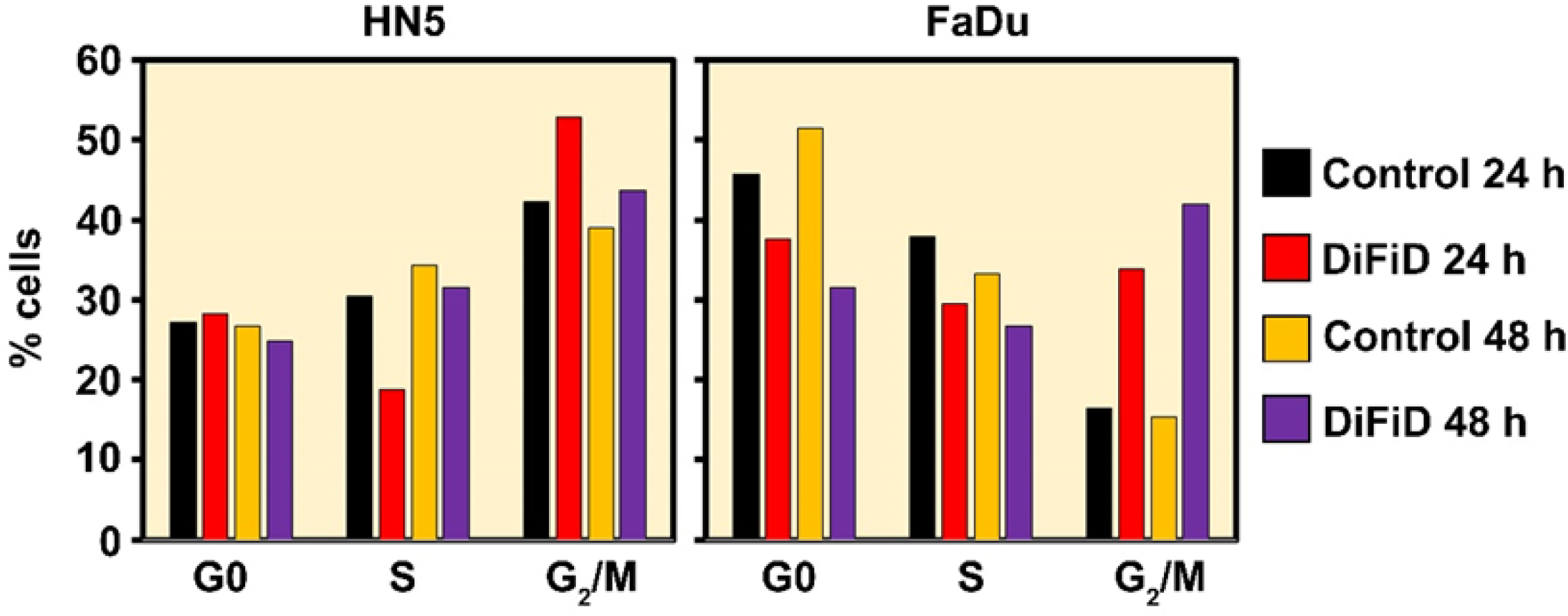
Quantification of cell cycle data from three independent experiments with HN5 and FaDu cells treated with DiFiD for 24 h or 48 h.

**Supplemental Figure 6.**
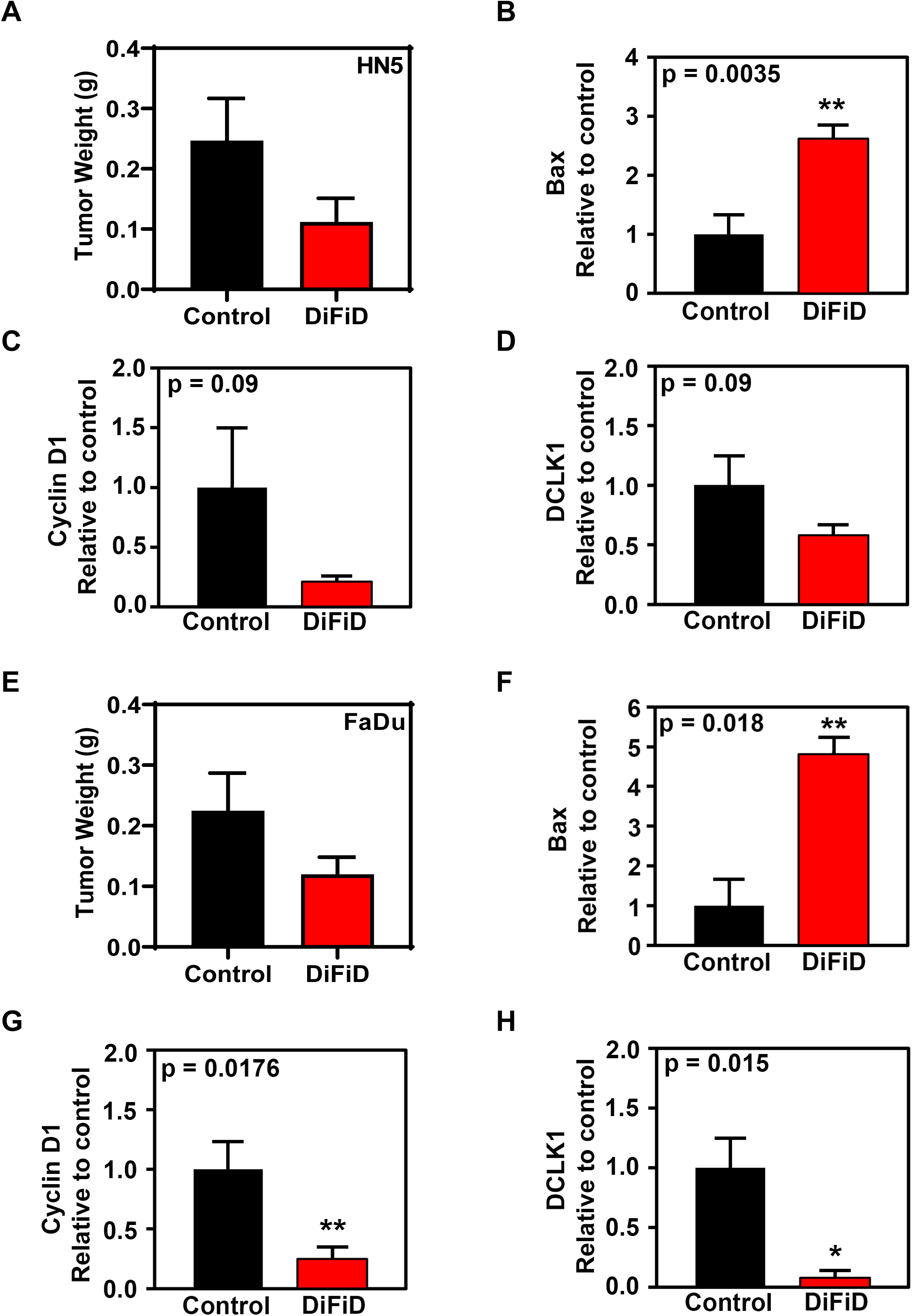
HN5 or FaDu (1 x 10^6^) cells were inoculated subcutaneously into the right flank of athymic *Foxn1^nu/nu^* mice. DiFiD (2 mg/kg) or vehicle control (DMSO) was administered by intraperitoneal injection once daily for 15 days. HN5 Tumor weight (A) (n=7 mice/group), are depicted. HN5 xenograft tumors were subjected to biomarker analyses. Densitometric analysis of HN5 xenograft tumors immunoblot signals for (B) Bax (C) Cyclin D1 (D) DCLK1. (E) FaDu tumor weight (n=10 mice/group). Densitometric analyses of FaDu tumor signals for (F) Bax (G) Cyclin D1 (H) DCLK1 are depicted graphically, including ± SEM, *p<0.05.

